# DNA Motors Powered by Exonuclease III for Autonomous Rolling Motion and Biosensing Applications

**DOI:** 10.1101/2025.11.26.690764

**Authors:** Yusha Imtiaz, Joshua Hardin, Bakai Sheyitov, Luona Zhang, Alexander Foote, Mohamed Husaini Bin Abdul Rahman, Krista Jackson, Selma Piranej, Alisina Bazrafshan, Khalid Salaita

## Abstract

Nucleic acid-based synthetic motors emulate key behaviors of biological machines, enabling applications in biosensing and nanoscale actuation. Among the reported synthetic motors, RNase H-powered motors offer high speed and processivity with demonstrated applications in computation and viral sensing. However, these motors rely on RNA as “fuel” source, limiting their stability. Here, we report the development of a robust, RNA-free DNA motor powered by Exonuclease III. These motors exhibit self-avoiding rolling motion driven by enzymatic hydrolysis of surface-bound DNA fuel strands, consistent with a burnt-bridge Brownian ratchet mechanism of translocation. We systematically optimized motor performance by chemically tuning the fluorescence reporter, DNA sequence composition, and surface fuel density. Fluorescence and brightfield microscopy revealed super-diffusive and Lévy-like stop-and-go dynamics under optimized conditions. Importantly, the established DNA-only architecture confers resistance to RNase degradation, and the system can be configured for motion-based biosensing via aptamer-functionalized components that selectively stall in response to viral targets. Beyond the significance of creating a chemically stable, tunable, and biosensing-compatible DNA motor platform, the work also establishes the modularity of the rolling motor platform and highlights how enzymatic diversity can expand their chemical and functional scope.

## Introduction

Living systems employ motor proteins to perform a variety of essential tasks ranging from cargo transport to muscle contraction and the dynamic re-arrangement of the cytoskeleton. These tasks are enabled by the activity of motor proteins, such as myosin, dynein, and kinesin that processively consume the chemical energy stored in adenosine triphosphate (ATP) to generate controlled mechanical motion^1,2^. Inspired by these motor proteins, researchers have strived to create synthetic motors and machines with the long-term potential applications in biosensing, computation, actuation, and robotics^3–10^. One prominent class of synthetic motors are based on “DNA walkers” which are nucleic acids engineered to move along one or two-dimensional DNA tracks and driven by different mechanisms including toe-hold mediated strand displacement, deoxyribozyme activity among others^4,8–10^. More recent designs of DNA walkers allow for autonomous movement, and have been employed for tasks such as signal amplification in biosensing and cargo transport at the nanoscale^4 9 11^. Nonetheless, DNA walkers still suffer from relatively limited speed and processivity in comparison to natural counterparts^5,11^, achieving only about 47 steps over a 12-hour period^12^.

To address the limited speed and processivity of DNA walkers, our group designed highly polyvalent, processive rolling motors driven by the RNase H enzyme^13–21^. These motors display micron/minute speeds, and their movement can persist for many hours. Rolling motors are comprised of DNA-modified spherical particles that can bind to complementary RNA fuel on planar surfaces. RNA-DNA heteroduplexes form at the motor-surface junction and become a unique substrate for RNase H. Thus, RNA hydrolysis exclusively occurs under the motor, creating a chemical gradient that drives forward motion of the motor to seek new Watson-Crick-Franklin base pairings. Rolling RNase H-based motors have been shown to be some of the fastest and most processive while demonstrating computation and sensing capabilities^15,18–20^. Importantly, the RNase H mechanism of translocation has been applied to virtually many types of motor chassis ranging from DNA origami nanostructures^15^ and gold nanoparticles^11,16^, to liposomes^10,22^ and micron-sized spheres^13–21^.

Nonetheless, the Achillies heel of RNase H-powered motors is the requirement for RNA fuel which is highly susceptible to degradation due to the ubiquity of RNases^19^. This is a major challenge when aiming to use motors for applications such as biosensing and computation that require highly robust responses in complex environments. To address this problem, here we report the development of a rolling motor that uses DNA fuel, which is orders of magnitude more stable than RNA. To drive translocation, we use Exonuclease III (ExoIII), a processive enzyme that hydrolyzes the 3’ blunt ends of DNA duplexes^23 24^. The enzyme releases one nucleotide at a time until the double stranded DNA (dsDNA) substrate becomes unstable and dissociates into single strands. The decision to employ ExoIII stems from its well-characterized multi-turnover reaction kinetics and the extensive control over its activity, allowing for fine-tuning of the motor’s speed and processivity^25–28^. By replacing RNA fuel with DNA, this system achieves robust and sustained operation under conditions that inhibit RNA-dependent designs. The ExoIII powered motors developed here maintain sustained, directional motion on DNA coated surfaces and routinely travel micron scale distances. By systematically tuning fluorophore identity, terminal sequence composition, and surface fuel density, we identified the molecular features that enhance motor speed, persistence, and overall stability. Detailed trajectory analyses reveal distinct bursts of movement that align with Lévy like stepping behavior previously observed in RNA based rolling systems^16^. To our knowledge, this represents the first DNA only rolling motor platform, and its fully modular design allows each component of the system to be independently tuned for performance, adaptability, and sensing applications.

## Results and Discussions

### Design & characterization of ExoIII-activated motors

The motor chassis is comprised of a 5 µm diameter silica particle modified with a dense monolayer of DNA oligonucleotides that we refer to as “DNA legs” (**Supplementary Table 1**). We achieved high-density DNA functionalization using a Cu-catalyzed click reaction (**Supplementary Fig. 1**, **Supplementary Fig. 2**). The DNA legs contained a 12-nucleotide domain complementary to the DNA monolayer on the chip surface (**Fig. 1a**). The chip surface was modified with a monolayer of fluorescently tagged DNA fuel complementary to the DNA legs (**Supplementary Fig. 3**). Upon introduction to the chip, the DNA motors settled and became immobilized by hybridizing to the fuel strands. We describe the rate of DNA leg-DNA fuel hybridization as ***k_on_***. Following the addition of ExoIII, the surface-bound DNA fuel is hydrolyzed in a step-wise manner from the 3′ end at a rate of ***k_cat_***, until the duplex becomes unstable and denatures at a rate of ***k_off_***. This sequence of binding, hydrolysis, dissociation, and re-binding is repeated, resulting in processive rolling motion. Because ExoIII activity irreversibly cleaves the previously visited foothold, motors move in a self-avoiding fashion through a mechanism consistent with the “burnt-bridge Brownian ratchet” model (**Fig. 1a**). As ExoIII typically digests both strands of a DNA duplex, we modified the DNA leg strands with a 3′ overhang of four thymidine bases^29^ and an inverted 3′-thymine modification to ensure hydrolysis occurred exclusively on DNA fuel. The motor trajectories were tracked in brightfield using time-lapse microscopy and subsequently analyzed with custom software and Python scripts, which is available on GitHub (**Supplementary Fig. 4**). **Figure 1b** shows a representative set of time-lapse images tracking a single motor for over 60 minutes using brightfield and fluorescence microscopy. The brightfield path illustrates the motor’s progression over time, and linescan analysis of this depletion track revealed a 21% decrease in fluorescein (FAM)-DNA fuel fluorescence intensity (**Fig. 1c**). This reduction serves as a local fuel depletion reporter, confirming that enzymatic digestion occurs along the motor’s path of mechanical displacement. On average, motors produced a 25% depletion in fluorescence intensity across tracks (**Supplementary Fig. 5**). Mean squared displacement (MSD) analysis of this trajectory yielded an alpha value (α) of 1.46, indicating super-diffusive motion (**Fig. 1d**). The alpha coefficient is obtained from the slope of the mean squared displacement (MSD) versus lag time (τ), following the relation _MSD (τ) ∝ τ_^α^. An α value of 1 corresponds to normal Brownian diffusion, α < 1 indicates sub-diffusive or constrained motion, α > 1 reflects super-diffusive transport driven by active processes, and α approaching 2 is characteristic of ballistic motion^30^. Additionally, we observed that a subset of dimerized motors exhibited ballistic motion as evidenced by linear depletion tracks and brightfield-tracked motion. In agreement with previous studies^13,15,16^, these anisotropic dimers displayed persistent unidirectional movement, providing direct support for a rolling mechanism underlying motor behavior. **Figure 1e** highlights three examples of such dimers: marginally interpenetrated, highly interpenetrating, and asymmetrical in size (left to right). The brightfield panels illustrate the distinct geometries, while the corresponding FAM channel reveals linear depletion tracks aligned with their paths, confirming ballistic displacement across all three variants. **Figure 1** highlights the motor behavior under what we have determined to be the most effective substrate sequence and assay conditions. Reaching this optimized state required extensive iteration, as ExoIII motor movement proved highly sensitive to a range of tunable parameters. Our efforts to establish a functional system and enhance motility began with modifications to the 3′ terminus of the DNA fuel, and the process of optimization is described in the following sections.

**Figure 1.**
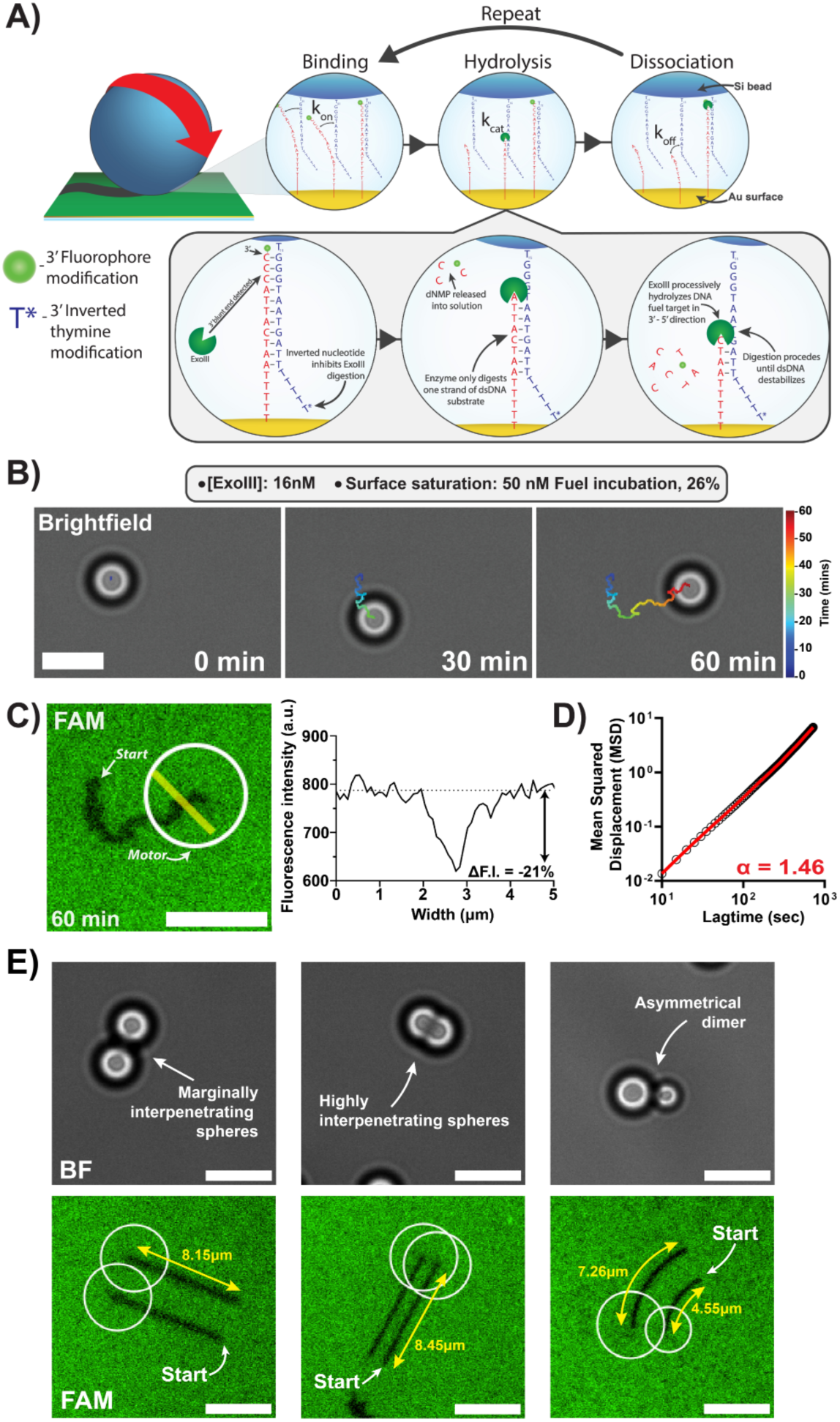
Design of Exonuclease III-powered rolling motors. **(A)** Top: Schematic for the hypothesized kinetic model to achieve rolling motion. The motors undergo displacement following the continuous cycle of binding, hydrolysis via ExoIII, dissociation, and re-binding of dissociated DNA legs with available fuel DNA substrates. Inset: ExoIII-mediated digestion of the fuel strand. An inverted 3′ nucleotide on the motor leg blocks cleavage, directing ExoIII to processively hydrolyze the surface-bound fuel strand in the 3′ to 5′ direction. As nucleotides are released and the duplex destabilizes, the motor leg dissociates and the binding–cleavage–rebinding cycle continues. **(B)** Real-time tracking of motor motion in brightfield (BF). Colored line represents trajectory of the motor obtained over 60 mins. Assay conditions: 16nM ExoIII, 50nM DNA fuel incubation in 1X rolling buffer. **(C) Left:** Representative image of motor (white circle) localization on the FAM-tagged DNA fuel surface after 60 mins of motion. **Center:** Background subtracted fluorescence depletion profile plot (right) from line scan (yellow line, left image) **(D)** MSD vs time plot of for the same motor in (B) and (C). The slope of the red line represents the alpha coefficient value (α = 1.46) **(E)** Anisotropic dimer particles moving in ballistic motion with varying levels of interpenetration (Left) Marginally interpenetrating dimer particle (Center) Highly interpenetrating dimer (Right) Asymmetrically sized dimer particle exhibiting preferential turning motion, where size imbalance biases the particle to curve rather than move in a straight ballistic trajectory. Assay conditions: 16nM ExoIII, 50nM DNA fuel incubation in 1X rolling buffer.

### Role of 3’ fluorophore modifications on motor behavior

To investigate the parameters that control the ExoIII-powered DNA motor system, we decided to use a standard set of conditions using 8 nM ExoIII, corresponding to 1 unit (U) of enzyme activity, and to incubate with 100 nM of DNA fuel, which was chosen to match conditions used in earlier rolling motor designs^13^. Under these baseline conditions, we sought to assess how 3’ terminus chemical modifications to the DNA fuel strand influenced motor behavior.

The 3’ end of the DNA fuel substrate serves as the initial site of hydrolysis by ExoIII, making it a critical determinant of motor performance^23,31^. Thus, chemical modifications at this position can alter both the effective enzyme *k*_cat_ and Michaelis-constant (*K_M_)*, thus influencing both overall motor speed and processivity^26^. Fluorophore labeling at the 3’ end enables real-time visualization of DNA fuel hydrolysis and its spatial localization with motor motion, allowing validation that enzymatic activity is driving particle translocation. We tested four groups: an unmodified strand and strands labelled with fluorescein (FAM), AlexaFluor488 (AF488), and Cyanine 3 (Cy3). These dyes differ in charge, hydrophobicity, and steric bulk at the 3′ end, giving us a way to test how the local chemical environment affects ExoIII activity and motor dynamics (**Fig. 2a**). Composite fluorescence images of motors following 30-minute rolling experiments with all three fluorophore-labeled DNA fuel variants showed the presence of small, circular depletions localized to the motor starting positions, which indicates ExoIII activity at the motor-fuel substrate complex (**Fig. 2b**). Through timelapse brightfield tracking, the coordinates (0,0) to demonstrate time-dependent outward motion of the motors (**Fig. 2c**). Utilizing the obtained trajectory data, we then calculated the net displacement, defined as the change in position between the initial and final timepoints (t = 30 min). This revealed that motors on unmodified DNA fuel exhibited the largest overall net displacement (mean = 1.67 ± 0.96 µm), although this condition did not permit depletion track visualization via fluorescence. Among the motors tested on fluorophore-modified DNA fuel, those using FAM strands exhibited the greatest motion, with a mean net displacement of 1.17 ± 0.60 µm. In comparison, motors tested on AF488 and Cy3 fuel surfaces produced smaller mean displacements of 0.85 ± 0.46 µm and 0.80 ± 0.52 µm, respectively (**Fig. 2d, left**). To evaluate the diffusional characteristics of motor trajectories, we calculated α-values. The mean α-values were 1.09 ± 0.21 for the unmodified fuel, 1.09 ± 0.16 for FAM, 1.06 ± 0.19 for AF488, and 0.93 ± 0.34 for Cy3. Note that alpha values are more error prone when net displacements are dampened, since localization error (83 ± 43 nm) from the tracking software^32^ can have a disproportionately larger effect on the calculation (**Supplementary Fig. 6**). As a result, α analysis is prone to noise for the slow moving or stalled motors (**Fig. 2d, right**). To better understand why 3’ dye conjugation to the DNA fuel impacted motor speed, we treated 100 nM DNA fuel-DNA leg duplexes (**Fig. 2e**) with 8 nM ExoIII for 15 min and resolved the products on a 10% native PAGE gel (**Fig. 2f**). In the nucleic acid stain channel, the ExoIII-treated unmodified DNA fuel showed a small but consistent increase in mobility, suggesting removal of one or a few 3′ terminal nucleotides, as the full band migrated lower than the untreated control. This is consistent with our calculations showing that after cleavage of a few nucleotides, the duplex becomes unstable, dehybridizes and no longer offers a viable substrate for the enzyme (**Supplementary Fig. 7**)^31,33^. Both FAM and AF488 bands showed significant enzyme-enabled digestion in contrast to Cy3, which resulted in minimal substrate depletion. We note that the FAM and AF488–modified substrates appear more concentrated with nucleic acid stain due to spectral overlap between the stain’s emission and the 488 nm channel. Hence, the intensity of the bands in nucleic acid stain channel was less useful for the dye-DNA conjugates. The fluorophore specific channel (488nm) showed drastic changes because single nucleotide cleavage results in loss of fluorescence for the DNA fuel substrate.

**Figure 2.**
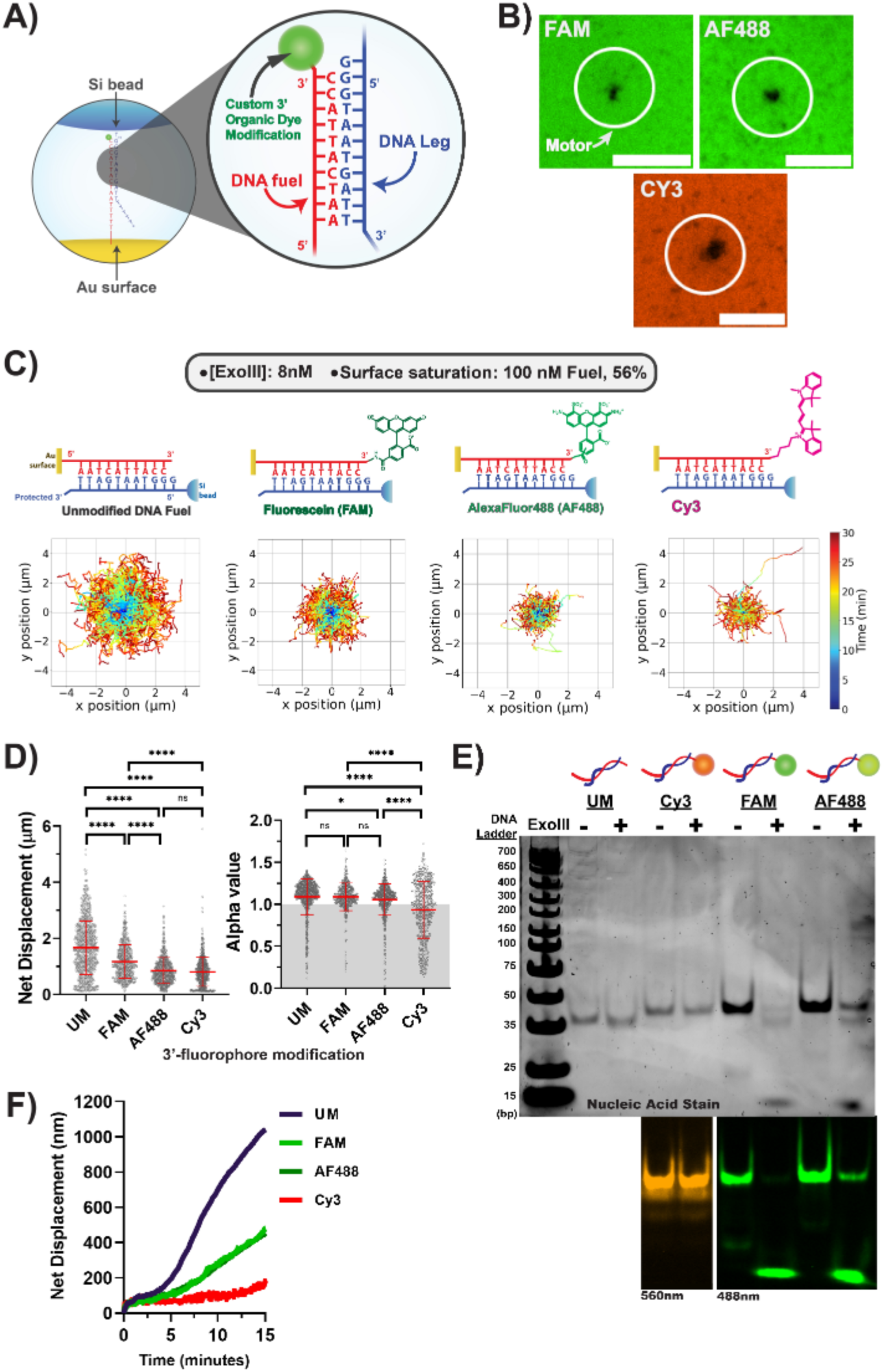
Influence of 3′ End Fluorophore Modifications on Motor Behavior. **(A)** Schematic of the motor system highlighting the tested sequence and the position of the 3’-fluorophore modifications. **(B)** Fluorescence images showing substrate depletion tracks after 30 minutes for FAM-, AF488-, and Cy3-labeled fuels. White circles denote motor bead localization. Scale bar = 5 µm. **(C)** Ensemble trajectory plots for motor motion for each fluorophore modification tested (left to right): unmodified, FAM, AlexaFluor488, Cy3. **(D)** Ensemble measurements for net displacement (left) and alpha values (right) from brightfield tracking of the motors for each fluorophore modification. The error bars represent the standard deviation of n = 700-900 motors. **(E)** 10% native PAGE gel showing ExoIII digestion of tested substrates. Unmodified -(UM), Cy3-, Fluorescein (FAM), and AlexaFluor488 (AF488)-labeled duplexes were incubated with (+) or without (–) 8 nM ExoIII. Nucleic acid stain and fluorescence imaging (560 nm, 488 nm). **(F)** Mean cumulative net displacement over time for each tested fluorophore modification.

The corresponding motor behavior was further assessed by analyzing the cumulative net displacement over a 15 min time window, matching the conditions used for gel analysis (**Figure 2F**). This analysis revealed that motors on unmodified DNA fuel exhibited the greatest sustained motion, while motors operating on FAM- and AF488-modified fuels showed similar cumulative net displacements. By contrast, motors on Cy3 fuel displayed minimal displacement, consistent with the gel results. The proposed mechanistic models of ExoIII activity suggest that the enzyme active site interacts with the 3’ nucleic acid substrate through base-stacking interactions with the key residues phenylalanine 213 and tryptophan 212^25^. Thus, we hypothesize that Cy3’s ability to engage in base-stacking interactions with DNA^34^ leads to inhibition of ExoIII activity, delaying substrate turnover. Surprisingly, we found that at later time points (>30 min), Exo III displayed greater non-specific 3’ ssDNA hydrolysis when presented with Cy3-DNA compared to FAM and AF488 ssDNA (**Supplementary Fig. 8**). Taken together, these results show that Cy3 is a poor reporter for ExoIII-powered motors due to non-specific substrate degradation and diminished specific activity. Accordingly, we identified FAM as a reliable fluorescence reporter for tracking motor depletion in the wake of moving particles.

### DNA fuel sequence significantly alters motor motility

We next explored how altering the sequence composition at the 3′ end of the DNA fuel affects motor motility. ExoIII differs fundamentally from RNase H not only in substrate preference but also in the mechanism by which it processes nucleic acid backbones. While RNase H is an endonuclease that cleaves the RNA strand within RNA–DNA hybrids at random positions, ExoIII functions as an exonuclease that binds to nicked or blunt 3′ termini of duplex DNA and hydrolyzes nucleotides in the 3′ to 5′ direction^27^.

This processive activity motivated the design of an asymmetric DNA fuel in which the 3’ region was enriched in cytosine bases. In this configuration, the substrate duplex forms with a high k_on_ and remains stable prior to enzymatic digestion. As ExoIII progressively removes terminal cytosine residues, the substrate stability decreases abruptly in a stepwise manner. NuPack simulations show that the substrate melting temperature (T_m_) drops by ∼8 ⁰C with each cytosine removed (**Supplementary Figure 7**). This design therefore increases k_off_ upon enzyme addition, promotes motor release, and allows the freed DNA leg strands to rehybridize with nearby undigested sites (**Fig. 3a**). When designing the asymmetric substrate, we anticipated a goldilocks-like relationship between DNA leg-DNA fuel stability and motor speed. Specifically, longer DNA leg-DNA fuel duplex would result in diminished motor activity as multiple cleavage events are needed to release the DNA leg and allow for motion; and conversely if the DNA substrate is unstable, then the motor would transition to being a Brownian diffuser which would prevent ExoIII activity.

**Figure 3.**
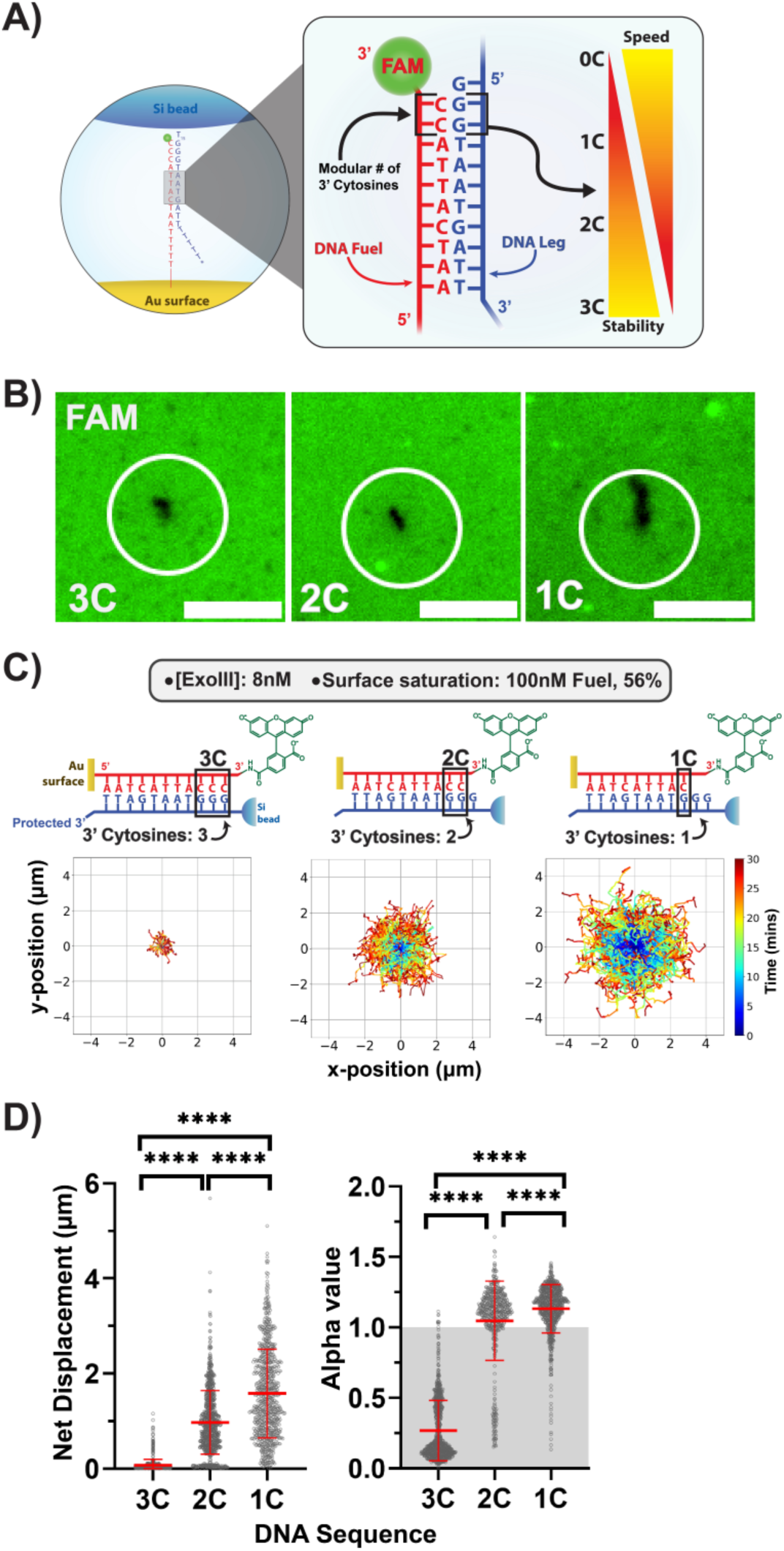
Decreasing Terminal Cytosines Boosts ExoIII Motor Performance. **(A)** Schematic showing variation in 3′ terminal cytosines on the DNA fuel and their expected impact on ExoIII activity and motor speed. **(B)** Fluorescence images of substrate depletion for motors on fuel strands with 3, 2, and 1 terminal 3’ cytosine(s). **(C)** Ensemble trajectory plots for motor motion using 3’-FAM modified DNA fuel with 3, 2, 1 terminal 3’ cytosine bases. Experiments were performed in 8nM ExoIII + 56% surface saturation assay conditions. **(D)** Ensemble measurements for net displacement (left) and alpha values (right) from brightfield tracking of the motors for each sequence variation. The error bars represent the standard deviation of n = 700-900 motors. All pairwise comparisons between conditions were statistically significant (p < 0.0001). Scale bars: 5 µm.

To evaluate this tradeoff, we designed three FAM-labeled DNA fuel variants containing three, two, or one cytosines at the 3′ terminus and monitored motor performance on each surface. Fluorescence imaging revealed small localized substrate depletion spots for 3C and 2C fuel conditions, consistent with previous experiments (**Fig. 3b**). Motors running on 1C fuel, however, produced noticeably longer tracks compared to those shown in **Figure 2**. The ensemble trajectory plots and net displacement analysis supported these findings with a clear trend in motor performance across the sequence variants (**Fig. 3c, d**). Motors on the 1C fuel exhibited the highest levels of movement with a mean displacement of 1.58 ± 0.93 μm. The 2C fuel resulted in motors with a mean net displacement of 0.97 ± 0.67 μm, and the 3C fuel with 0.07 ± 0.12 μm. These results support the idea that reducing the number of terminal cytosines increases the dissociation rate of DNA fuel following ExoIII digestion, leading to faster turnover and enhanced displacement. A similar trend was observed in α-values. Motors on the 1C and 2C fuels demonstrated predominantly super-diffusive behavior, with α-values of 1.13 ± 0.17 and 1.05 ± 0.28, respectively (**Fig. 3d**). In contrast, the 3C condition yielded a significantly lower mean α of 0.27 ± 0.21 and is more prone to the influence of localization error. These results confirmed the positive impact of reduced cytosine content on motor dynamics. We also tested a 0C design to further destabilize the DNA fuel substrate, but it consistently failed to produce visible depletion tracks (**Supplementary Fig. 9**). Thus, we concluded this substrate condition to result in Brownian diffusers rather than active motors. Among all tested designs, the 1C condition provided the most favorable balance at this DNA fuel surface saturation as it supported strong displacement, robust directionality, and consistently visible depletion tracks. These results also highlight how sensitive the system is: a difference of only three hydrogen bonds at the duplex terminus can determine whether motors stall, idle often, or move processively. The 1C design therefore represents the goldilocks sequence that offers just enough stability to promote binding while still allowing efficient release after ExoIII digestion, making it the most suitable fuel configuration under the conditions tested..

### DNA fuel density regulates motor speed and diffusion behavior

Having screened the DNA fuel sequence design space, we next investigated how both the surface density of DNA fuel and enzyme concentration influence motor performance. To vary the DNA fuel density, FAM-labeled DNA fuel was incubated at concentrations ranging from 25 nM up to 500 nM for 6 hours to hybridize with complementary anchor strands (**Fig. 4a**). After rinsing excess oligo, the relative surface density of DNA fuel was quantified using FAM fluorescence intensity. Representative images confirmed progressive increases in surface coverage across this range (**Fig. 4b**). Triplicate measurements and fitting of the data produced a calibration indicating that 25 nM, 50 nM, 75 nM, 100 nM, 250 nM, and 500 nM incubation concentrations corresponded to 4%, 20%, 43%, 56%, 90%, and 97% surface saturation, respectively (**Fig. 4c**). While we are not quantifying the absolute surface density in these measurements, note that our prior calibration showed that 100 nM incubation of RNA fuel strands results in a density of 5× 10^4^ oligonucleotides/micron^13^. We next measured motor performance using these surfaces. At 56% fuel saturation, motors driven by 8 nM ExoIII exhibited limited displacement and compact trajectories (mean net displacement = 0.97 ± 0.67 μm), whereas at 20% saturation, the motors using the same enzyme concentration produced more extended motion (mean net displacement = 1.65 ± 1.04 μm). This suggested that the 20% surface coverage produced improved net displacements (**Fig. 4d**). We next increased the enzyme concentration to 16 nM which further enhanced mean net displacement to 2.73 ± 1.49 μm (**Fig. 4d**). We observed that the mean α-values also supported this trend with the 16 nM ExoIII at 20% saturation condition having the greatest α-value of 1.24 ± 0.20, followed by those of 8 nM ExoIII at 20% at 1.17 ± 0.16, and then finally 8 nM ExoIII at 56% saturation at 1.05 ± 0.28 (**Fig. 4e**).

**Figure 4.**
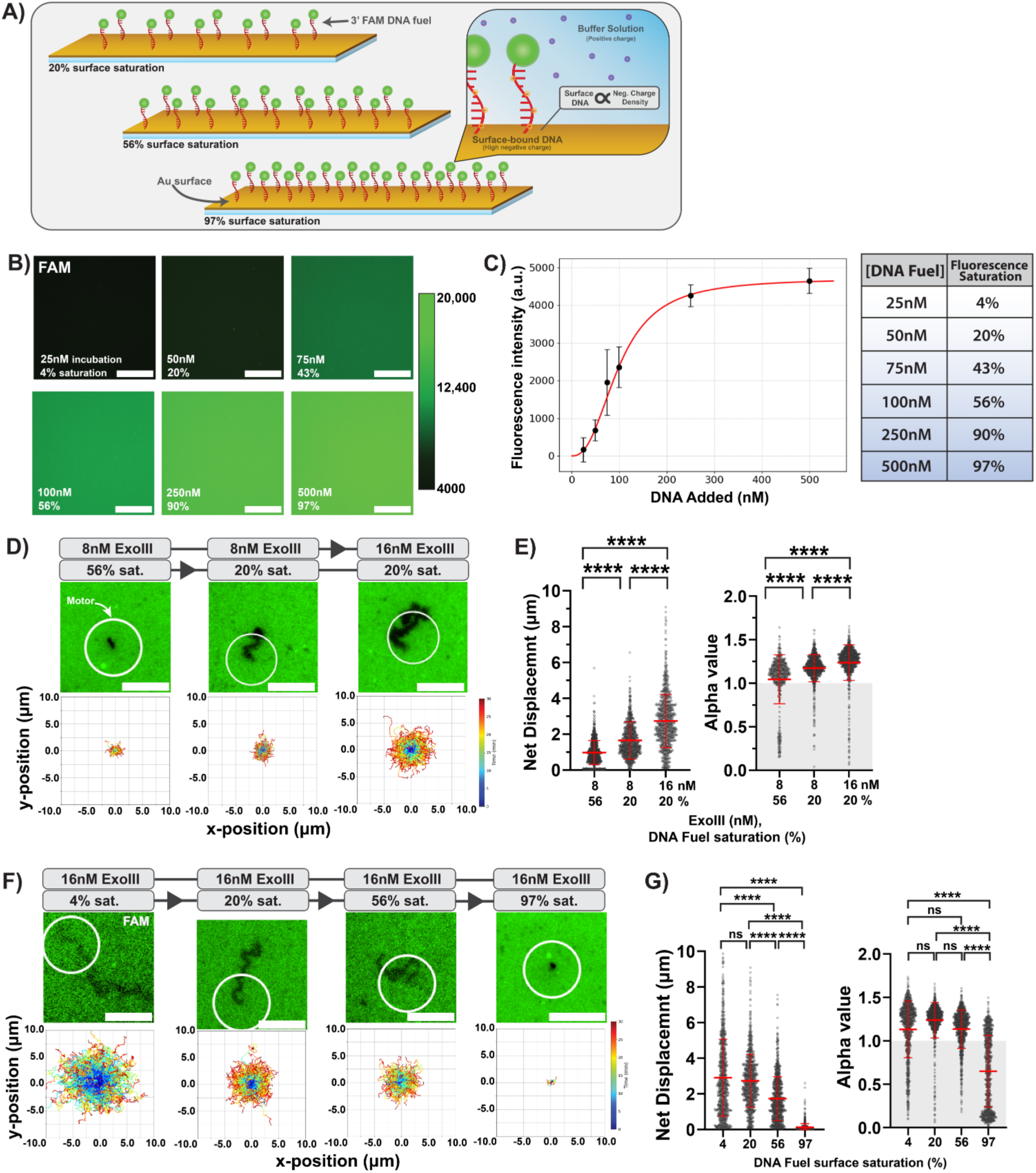
DNA Fuel Surface Density Regulates ExoIII Motor Performance. **(A)** Schematic illustrating increasing surface coverage of 3′-FAM–labeled DNA fuel strands on the gold surface, from sub-saturating (20%) to full saturation (78%). Inset: increased surface DNA density enhances negative surface charge. **(B)** Fluorescence images surfaces incubated with varying concentrations of 2C FAM DNA Fuel. Scale bars: 100 µm. **(C)** Fluorescence saturation curve of surface-bound FAM-labeled DNA fuel as a function of fuel concentration added. **(D)** Top: Fluorescence images of motor depletion tracks after 30 minutes of rolling on surfaces prepared with 8nM ExoIII + 56% saturation, 8nM ExoIII + 20% saturation, and 16nM ExoIII + 20% saturation, respectively. Bottom: Ensemble trajectory plots of representative single-motor trajectories with the same assay conditions. Scale bars: 5 µm. **(E)** Ensemble measurements for net displacement (left) and alpha values (right) from brightfield tracking of the motors across same assay conditions presented in (D). The error bars represent the standard deviation of n = 700-900 motors. **(F)** Top: Fluorescence images of motor depletion tracks after 30 minutes of rolling on surfaces prepared with 25, 50, 100, and 500 nM DNA fuel, corresponding to 5%, 20%, 56%, and 97% saturation, respectively. Bottom: Ensemble trajectory plots of representative single-motor trajectories across surface saturations. **(G)** Ensemble measurements for net displacement (left) and alpha values (right) from brightfield tracking of the motors

We next systematically tested motor performance under varied surface saturation using the 16 nM enzyme concentration. Motors on 4% and 20% surfaces exhibited the longest trajectories and greatest net displacements, with mean values of 2.91 ± 2.16 µm and 2.73 ± 1.49 µm, respectively (**Fig. 4f–g**). In comparison, the two higher fuel saturations demonstrated decreased motility with the 56% fuel saturation resulting in a mean displacement of 1.74 ± 1.23 µm, and the motors at 97% fuel saturation producing only short trajectories at a mean displacement of 0.13 ± 0.21 µm. Interestingly, we observed that motors running on the 20% saturation surfaces produced the highest mean alpha values of 1.24 ± 0.20. Motors running on the 4% and 56% saturated surfaces had similar, but lower mean α-values of 1.13 ± 0.33 and 1.14 ± 0.22, respectively. Finally, motors on the 97% saturated surface produced a α-value of 0.65 ± 0.41 (**Fig. 4g**). The 4% condition contained a subset of motors with greater α-values than those observed at 20%, but the overall mean was reduced because this condition also contained a larger proportion of motors exhibiting sub-diffusive behavior.

One plausible explanation is that at low fuel densities, particles mediate the rapid depletion of the few fuel strands they bound to, and with no nearby substrates available for rehybridization, remain immobilized.

Efficient translocation therefore requires a critical density of DNA fuel and DNA legs that supports processive forward motion, and densities above this threshold begin to impede that progression. This threshold density is dependent on enzyme concentration, with higher enzyme levels requiring a correspondingly higher fuel density. Importantly, the threshold for processive motion also depends on the stability of individual DNA fuel-leg duplexes. This was confirmed when we tested the 1C and 0C DNA fuel variants at lower surface saturations. In these measurements we did not observe enhancement in motility and trajectories remained limited (**Supplementary Fig. 10**). Together, these results demonstrate that motor performance is maximized at intermediate DNA fuel surface densities (20% of maximum saturation) when using the 2C design, highlighting the importance of balancing substrate stability with enzymatic turnover and k_on_ rate.

### In-depth analysis of motor behavior

Having established an RNA-free DNA rolling motor, we next characterized motor behavior in greater depth at 20% DNA fuel saturation. In many of our experiments, we observed that ExoIII-powered motors frequently exhibited bursts of speed. This behavior is clearly visualized in a representative single-motor trajectory plot over time and the corresponding instantaneous speed profile (**Fig. 5a left, center**). Sharp fluctuations in speed can in ensemble trajectory plots and corresponding speed-resolved plots (**Fig. 5b left, center**). However, when the instantaneous speeds were averaged across the entire ensemble, we found that this type of rendering obscured individual trajectories that displayed speed fluctuations and dynamic bursts (**Fig. 5b right**). We suspected that this behavior was consistent with the Lévy walk classification^35^, a form of anomalous diffusion characterized by intermittent bursts of long steps separated by pauses or short exploratory movements, producing a heavy tailed step size distribution. To test this, we fitted step size distributions to a power law model, *P* (*x*) ∝ *x*^− *μ*^, where ^*x*^ denotes motor step size and µ is the Lévy exponent (**Fig. 5c**). Both enzyme concentrations produced heavy tailed distributions characteristic of Lévy walks (1 < μ < 3), with μ exponents of 1.94 ± 0.02 at 8 nM and 2.38 ± 0.01 at 16 nM. The lower μ at 8 nM ExoIII reflects a heavier tailed distribution with more rare long steps, whereas the higher μ at 16 nM ExoIII indicates more frequent bursts of intermediate to large displacements.

**Figure 5.**
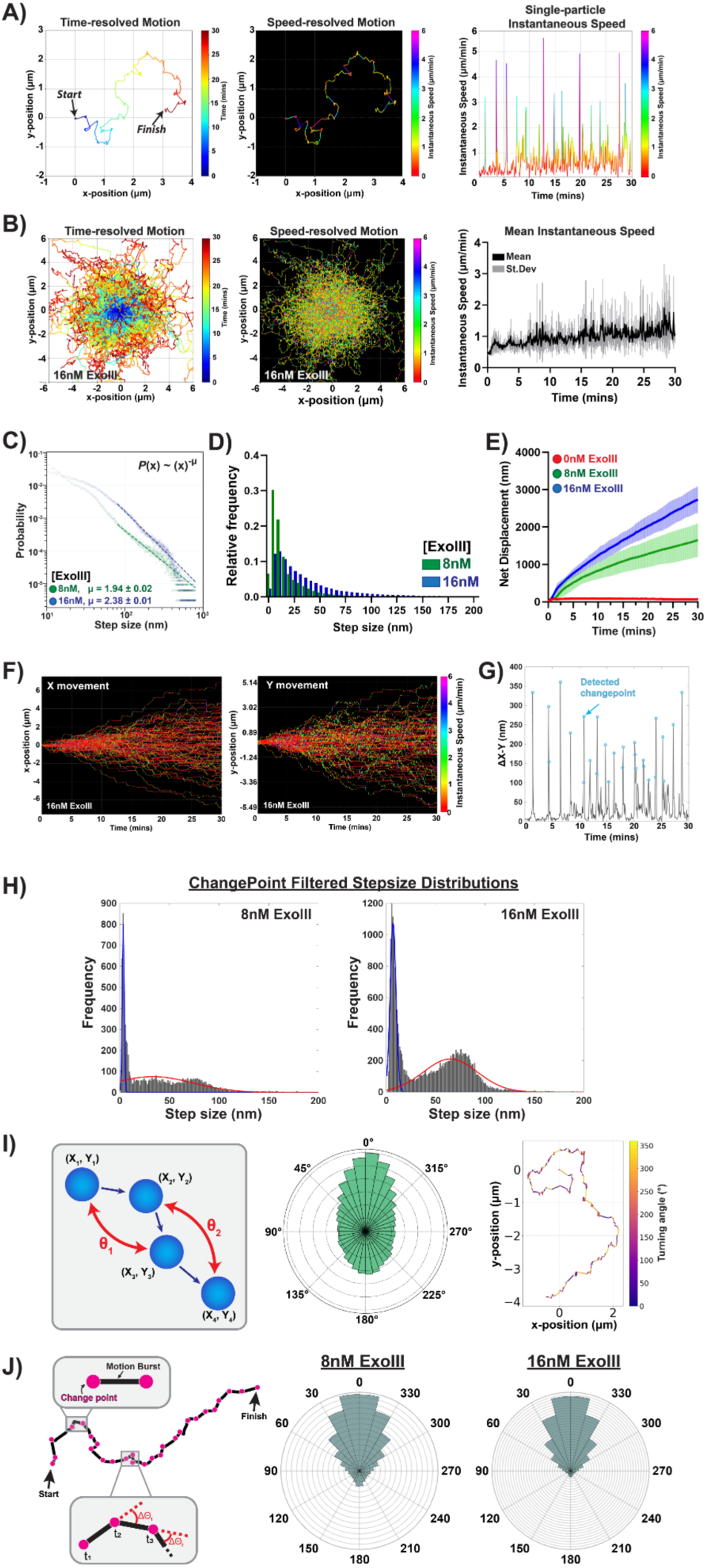
Single-Particle Analysis of Motor Behavior. All motors were run in optimized conditions:16 nM ExoIII + 20% surface saturation. **(A)** Left: Time resolved trajectory plot of a single motor with optimized conditions. Center: Speed-resolved trajectory plot. Right: Speed trace over timelapse for single motor with instantaneous speed represented by line color. **(B)** Left: Time resolved trajectory plot of motor motion with optimized conditions. Center: Speed-resolved trajectory plot. Right: Time-dependent mean (dark grey) instantaneous speed of n = 900 motors with standard deviation (light grey). **(C)** Power-law fits of ExoIII motor step lengths with varying ExoIII concentrations obtained by computing exponential fits of step size probability distributions. Motors with 1 < µ < 3 are classified as to demonstrate Levy-type motion. **(D)** Distribution of motor step sizes Green: 8nM, blue: 16nM. **(E)** Cumulative net displacement over time of motors with varying ExoIII concentrations (red: 0nM, green: 8nM, and blue: 16nM). **(F)** XY axes separated motion of motors. Plot contains n = 100 motors for visual clarity. **(G)** Δ XY of motors with detected changepoints circled in cyan. Changepoint algorithm detects points of high speed motion relative in motor trajectories relative to the entire dataset. **(H)** Distributions of motor change-point speeds comparing 8nM & 16nM ExoIII conditions. **(I)** Left: Scheme depicting turning angles of motors between consecutive trajectory points. Center: Distribution of turning angles on a polar plot. Right: Turning angles overlayed on a motor trajectory with the colormap representing angle values. **(J)** Left: Scheme of changepoints overlayed with a 2D trajectory plot. Directionality of motors is calculated by

To investigate how enzyme concentration modulates the dynamics of the Lévy walk speed bursts, we first analyzed the motor trajectories without any data filtering to examine how step-size distributions change with increasing ExoIII concentration. As the ExoIII concentration increased from 8 nM to 16 nM, we observed a larger population of long steps (**Fig. 5d**), consistent with greater overall motility reflected in the cumulative mean net displacement over time (**Fig. 5e**). Although ensemble measurements reveal higher motility at elevated enzyme concentrations, they obscure the brief acceleration and pausing events that govern individual motor motion.

To resolve these finer dynamics, we next performed single-particle analyses to isolate and examine active periods within each trajectory. Individual trajectories were visualized by plotting motor displacements along the x and y axes over time (**Fig. 5f**). After smoothing, the frame-to-frame derivative of displacement (ΔXY) was computed to reveal distinct bursts and pauses in motion (**Fig. 5g**). Sharp increases in ΔXY marked motility events were automatically detected using a changepoint algorithm, which identifies statistically significant shifts in the mean or variance of the signal to delineate active and stationary phases within each trajectory (**Supplementary Fig. 11**). Finally, applying the changepoint algorithm revealed that while motors at both enzyme concentrations achieved similar maximum step sizes, higher ExoIII concentration increased the frequency of large displacement bursts. In this system, Lévy like motion arises because motors remain stationary until all footholds beneath them the enzyme concentration shortens these waiting periods by accelerating substrate digestion, producing more frequent bursts and greater overall motility (**Fig. 5h**).

We next investigated whether these bursts of motion occurred with any directional preference. To quantify this, we calculated the turning angle (Δθ) between successive displacement vectors along each motor trajectory, defined by the positional coordinates of consecutive frames (*x*_1_, *y*_1_), (*x*_2_, *y*_2_), (*x*_3_, *y*_3_) (**Fig. 5i, left**). The angle between each pair of steps represents the degree of directional change, where small Δθ values indicate forward persistence and larger Δθ values reflect more random or reversing motion. When all trajectory data were analyzed, the polar plots showed a weak preference for forward motion, with most motors displaying angular changes within ±30° (**Fig. 5i, center**). However, this trend is partly masked by localization noise within stationary periods, as many angular data points originate from frames with minimal movement, as seen in the trajectory overlays with turning angles (**Fig. 5i, right**). To resolve this, we overlaid the previously identified change-points onto each trajectory to isolate the sections of active motion. By recalculating Δθ between change point positions, we obtained analysis of directionality during speed bursts (**Fig. 5j left**). The filtered distributions revealed the same overall forward bias but with reduced noise, confirming that directional persistence becomes more evident once nonmotile periods are excluded. Moreover, increasing the ExoIII concentration increased both the frequency and consistency of forwardly directed steps, consistent with enhanced processive motion at higher enzyme activity levels (**Fig. 5j, center & right**).

Together, these analyses establish that ExoIII-powered DNA motors move through bursts of directed, Lévy-like displacements that become more frequent and persistent with increasing enzyme concentration. Ultimately, our findings indicate that the behavior of the ExoIII motors is consistent with the mechanistic models described in previous work^14,16,17,36^, involving cyclic phases of substrate digestion, release, and rebinding (**Supplementary Fig. 12**).

### RNase resistance establishes DNA-only motors as a platform for biosensing

Thus far, RNA-fueled motors have shown the greatest speed, endurance, and processivity which has opened the door for applications in viral diagnostics and computation. A major challenge for RNA fueled rolling motors relates to their vulnerability to RNases. Accordingly, RNA fueled rolling motors have limited utility in the vast majority of biological samples^19^. To examine whether the current ExoIII-catalyzed system could overcome this limitation, we repeated our assay using the 2C-FAM fuel at 56% saturation. This higher density was chosen to heighten the sensitivity of the assay, since crowded duplex layers reveal even weak RNase H activity and place additional constraints on rolling, enabling a clear evaluation of RNase H resistance and ExoIII dependence. To begin, we let the motors sediment and bind to the surface and then added 5 U of RNase H and monitored FAM surface density and particle positions (**Fig. 6a**). In the presence of 5 U RNase H alone, no visible motion or substrate degradation was observed over a 30 min window, consistent with the enzyme’s inability to hydrolyze DNA–DNA duplexes and ssDNA. However, when 16 nM ExoIII was added to the same RNase H–treated surface, we observed the appearance of distinct depletion tracks spatially localized with motors on brightfield, indicating substrate digestion and active rolling motion (**Fig. 6b**). Quantitatively, motors incubated with RNase H alone exhibited a mean net displacement of 0.12 ± 0.15 µm. A substantial portion of the motor population remained effectively immobile, displaying apparent motion comparable to the localization error of the ExoIII system. (**Supplementary Figure 6**). In contrast, the addition of ExoIII to the same RNase H–treated surfaces resulted in clear motor motion with a mean displacement of 1.35 ± 0.70 µm (**Fig. 6c, d**).

**Figure 6.**
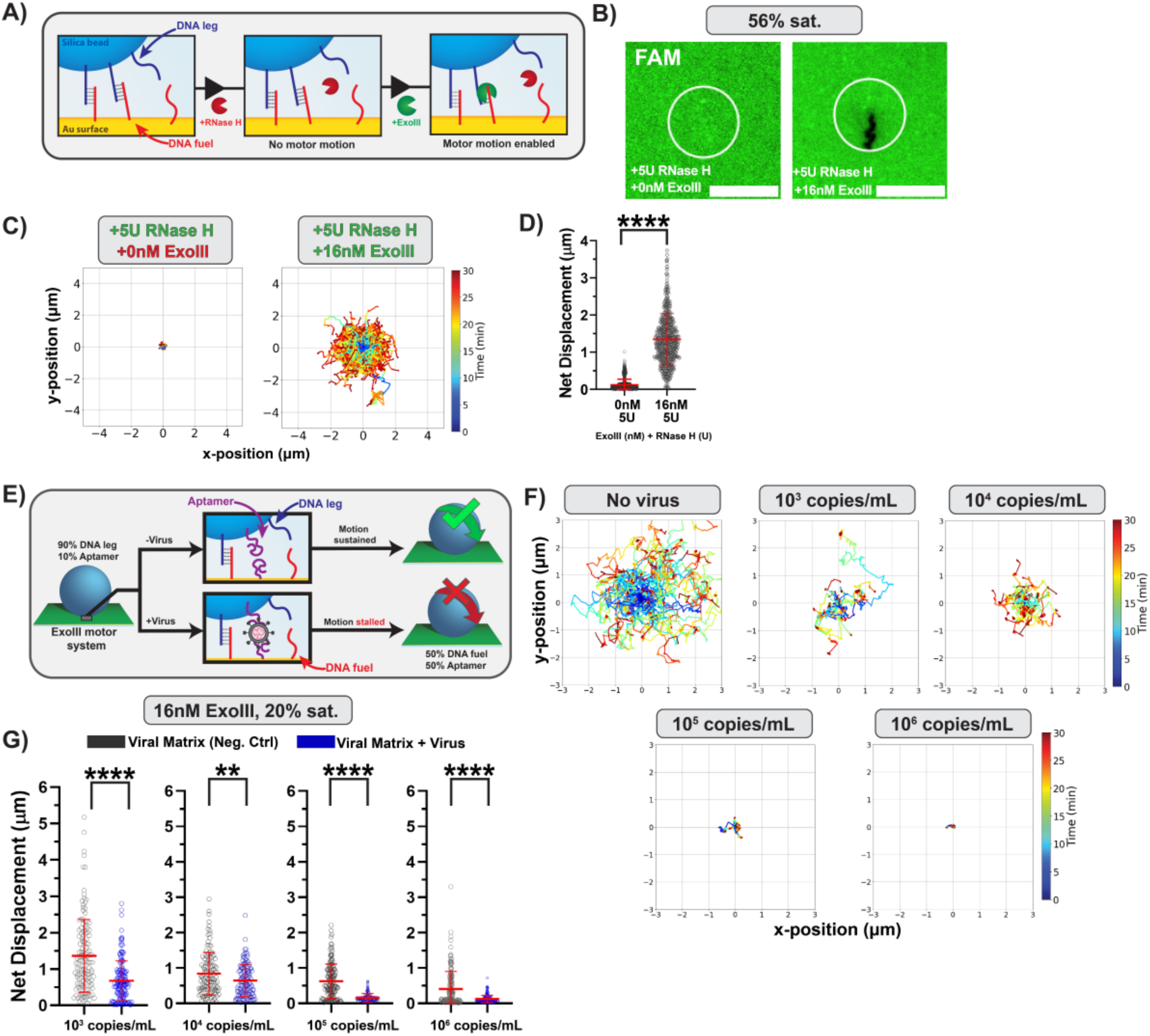
RNase Resistance and Biosensing Potential of ExoIII Motors. **(A)** Schematic illustrating the ExoIII motor system’s resistance to RNase degradation due to the DNA-based architecture. Motor activity occurs exclusively in the presence of ExoIII, enabling selective activation. **(B)** Fluorescence images of motor depletion tracks after 30 minutes of rolling with RNase H (left) and both RNase H & ExoIII (right). **(C)** Ensemble trajectory plots of representative single-motor trajectories with or without RNase H & ExoIII. **(D)** Ensemble measurements for net displacement (left) and alpha values (right) from brightfield tracking of the motors with or without ExoIII in the presence of RNase H. **(E)** Schematic demonstrating biosensing mechanism for the ExoIII system. A portion of the DNA leg & DNA fuel populations are substituted with aptamers specific for the viral target. The motors will move normally in the absence of the viral target but will consistently stall in its presence due to the formation of an aptamer-ligand binding interaction between both the motor-bead and surface. **(F)** Ensemble trajectory plots of representative single-motor trajectories tested with different viral target concentrations. **(G)** Ensemble measurements for net displacement (left) and alpha values (right) from brightfield tracking of the motors tested across viral target concentrations. Grey = motors + viral matrix - virus; Blue = motors + viral matrix + virus.

Alpha values for these experiments also result in the same trend (**Supplementary Fig. 13**). Together, these results confirm that ExoIII activity is necessary for motor function and, importantly, that the DNA-only design remains resistant to RNase H, enabling reliable operation in environments that would otherwise confound RNA-based motors.

Next, we tested if the ExoIII motor system could be used as a motion-based viral biosensor. To achieve this, we adapted the design of our previously developed RoloSense assay^19^, in which a fraction of the DNA sequences on both the motor chassis (10%) and the surface (50%) were replaced with a virus-specific aptamer (**Fig. 6e**). In this configuration, ExoIII motors exhibit continuous motion in the absence of viral targets but stall upon their introduction. The halt in motion occurs because the viral particles display multivalent antigens and are thus “sandwiched” by bridging the aptamer-modified strands on the surface with those on the motor chassis (**Fig. 6e**). For this work, we used the “SP6.34” aptamer reported by Schmitz et al.^37^ and incorporated a 3′-inverted thymine modification to protect it from ExoIII digestion. This aptamer is reported to have an approximate K_D_ of 13 nM against the spike protein of SARS-CoV-2^19^. The aptamers on both the motors and surface were Motors were first incubated with the viral samples for 30 min in viral matrix. Then, the motors were added to the surface and allowed to sediment for another 30 min. Finally, 16 nM ExoIII was then added to the solution and brightfield imaging was used to track particle motion. We measured motor dynamics with a range of increasing viral copies per sample of SARS-CoV-2 delta strain (B.1.617.2) (**Fig. 6f**). For samples incubated with 10^3^ copies/mL of SARS-CoV-2 the stoichiometry of motors to virions was 1:1. At this concentration, motors displayed the greatest mean displacement (0.68 ± 0.55 µm). At 10^4^ copies/mL (motors:virions 1:10) motors moved with a similar mean net displacement of 0.65 ± 0.45 µm (**Fig. 6g)**. At higher viral concentrations of 10^5^ (motor:virion = 1:100) and 10^6^ copies/mL (motor:virion = 1:200), however, we found that the motors stalled significantly with mean net displacements of 0.16 ± 0.20 and 0.13 ± 0.20, respectively. This progressive decrease in motility at higher viral concentrations is attributed to increased multivalent binding between the aptamer-functionalized motors, surface, and virions. As more binding sites become occupied, these crosslinking interactions restrict the motors’ ability to detach and rebind to new sites, resulting in more frequent tethering and pronounced sub-diffusive stalling behavior (**Supplementary Fig. 14**). Interestingly, the negative control samples, in which motors were incubated with viral matrix corresponding to the amount present in their respective virus sample volumes but without virions, also showed a gradual decrease in mean net displacement with increasing sample concentration. The viral matrix includes serum proteins and other potential DNA-binding biomolecules that likely stall motor through non-specific interactions. This effect is further supported by comparison to a “non-matrix” control, in which motors were incubated with 1× PBS according to the standard non-biosensing protocol (**Supplementary Fig. 15**). Together, these results demonstrate that the ExoIII motor system can serve as a motion-based biosensor, where motor stalling provides a direct and quantifiable readout of target recognition through aptamer-mediated binding. While the current implementation remains unoptimized, its performance establishes a strong foundation for future development of more sensitive and selective motor-based biosensors.

## Conclusion

We report the development of an RNA-free rolling DNA motor system powered by ExoIII. By systematically varying 3′ fluorophore modifications, terminal sequence composition, and surface fuel density, we identified the conditions that maximize motor displacement, directionality, and processivity. FAM emerged as the most effective fluorophore modification, reduced cytosine content improved substrate turnover, and intermediate surface densities provided the optimal balance between substrate stability and enzymatic accessibility. Single particle analyses revealed that ExoIII motors move through intermittent bursts of activity consistent with Lévy like dynamics, an observation aligned with our previous findings for RNase H powered motors, where higher enzyme concentrations produced more frequent bursts, greater directional persistence, and higher net displacements.

Importantly, the DNA only design confers resistance to RNase H degradation and supports motion based biosensing through aptamer functionalized substrates. In this assay, the presence of SARS-CoV-2 virions caused motor stalling due to multivalent aptamer binding between the motor, surface, and viral targets.

Negative controls containing equivalent amounts of viral matrix but no virions showed a gradual decrease in motility with increasing matrix concentration, demonstrating that the assay is affected by the presence of external media. Although the current biosensing implementation remains unoptimized, it demonstrates the feasibility of coupling ExoIII driven motion to molecular recognition events. Future optimization of surface passivation using PEG^38^, zwitterionic groups^39^, or other surfactants will likely lead to suppression of non-specific matrix interactions.

By expanding the enzyme-substrate scope of DNA motors, this work lays the foundation for creating new types of motors powered through other types of enzymes. We anticipate new possibilities in medical diagnostics, nanoscale robotics, and sensors.

## Methods & Materials

All chemicals were purchased from Milipore-Sigma unless noted otherwise. Stock solutions were made using Nanopure water (Barnstead Nanopure system, resistivity = 18.2 MΩ) herein referred to as DI water. Oligonucleotides were custom synthesized by Integrated DNA Technologies (Coralville, IA) and stored at 4⁰C. Oligonucleotide sequences and functional groups are all shown in Supplementary Table S1. The 5 µm aminated silica beads were obtained commercially by purchasing from Bangs Laboratory (SA06N). Exonuclease III was obtained from New England Biolabs (M0206L). UV-inactivated SARS-CoV-2 Isolate hCoV-19/USA/PHC658/2021 (Lineage B.1.617.2; Delta Variant), NR-55611, was contributed by Dr Richard Webby and Dr. Anami Patel. Thin Au films were generated using a home-built thermal evaporator system. All motor translocation measurements were performed in ibidi sticky-Slide VI0.4 17 × 3.8 × 0.4 mm^3^ channels on ibid #1.5H glass coverslips (10812).

### Microscopy

Bright-field and fluorescence images were acquired on a fully automated Eclipse Ti2-E Nikon Inverted Research Microscope using the Elements software package (Nikon), an automated scanning stage, a 1.49 NA CFI Apo TIRF ×100 objective, a 0.50 NA CFI60 Plan Fluor ×20 objective, a Prime 95B 25 mm scientific complementary metal oxide semiconductor (sCMOS) camera for image capture at 16-bit depth, a SOLA SE II 365 Light Engine as a solid-state white-light excitation source and a perfect focus system used to minimize drift during time-lapse. Fluorescence images of cyanine dyes Cy3, AlexaFluor488 and fluorescein dye FAM were collected using a TRITC filter set (Chroma, 96321) and EGFP/FITC/Cy2/Alexa Fluor 488 filter set (Chroma, 96226). Images were initially taken with an exposure time of 100 ms and then once more at 500ms, with the former being used for analysis and the latter for use as representative images. All imaging was conducted at room temperature.

### Thermal evaporation of gold films

A #1.5H glass coverslip (Ibidi, 10812) was cleaned by sonication in 200 proof ethanol (Fisher Scientific, 04-355-223) for at least 10 minutes. The cleaned coverslip was then dried in an oven for 15 minutes and left out on a benchtop to return to room temperature. The clean, dried coverslip was then mounted in a home-built thermal evaporator chamber in which the pressure was reduced to 50 × 10−3 torr. The chamber was purged with N_2_ three times and the pressure was reduced to (1–2) × 10−7 torr using a turbo pump with a liquid N_2_ trap. Once the desired pressure had been achieved, a 1.2-nm film of Cr was deposited onto the slide at a rate of 0.1 Å s–1, which was determined by a quartz crystal microbalance. After the Cr adhesive layer had been deposited, 3-nm layer of Au was deposited at a rate of 0.2 Å s–1. The Au-coated samples were used within 1 week of deposition.

### Fabrication of DNA monolayers

An ibidi sticky-Slide VI^0.4^ flow chamber (ibidi, 80608) was adhered to the Au-coated slide to produce six channels (17 × 3.8 × 0.4 mm^3^). Next, thiol-modified DNA anchor strands were added to each of the channels using a 50 μl solution of 1 μM DNA anchor in 1 M KHPO_4_ buffer. The gold film was sealed in a large plastic petri dish with a microtube filled with DI water to maximize localized humidity. The gold film in the dish was then sealed with Parafilm to further prevent evaporation, with the reaction taking place overnight at room temperature. After incubation, excess DNA was removed from the channels by rinsing with ∼5 ml DI water. To block any bare gold sites, which non-specifically interact with nucleases, and maximize the hybridization of RNA to the DNA anchoring strands, the surface was passivated by backfilling it with 100 μl of a 100 μM solution of 11-(mercaptoundecyl)hexa(ethylene glycol) (SH-PEG; Sigma-Aldrich, 675105) solution in ethanol for 6 h. Excess SH-PEG was removed by rinsing with ∼5 ml DI water. Finally, 100 μl of 100nM DNA fuel strands (in 1X PBS) with partial complementarity with the DNA anchor strand were added and left to incubate for 6 hr. The gold film was again placed in a large petri dish and sealed with Parafilm to prevent evaporation. The final DNA monolayer remained stable for several days.

### Synthesis of azide-functionalized silica beads

Before functionalization with azide, 2mg of aminated silica beads (Bangs Laboratory, SA06N) were washed by centrifugation for 5 min at 15,000 r.p.m to remove any impurities. The supernatant was discarded, and the resulting particles were resuspended in 1 ml DI water. This was repeated three times and the supernatant was discarded after the final wash. The azide-functionalized particles were then synthesized by mixing 2 mg aminated silica beads with 2 mg azidoacetic *N*-hydroxysuccinimide ester (BroadPharm, BP-22467). This mixture was subsequently diluted in 100 μl dimethylsulfoxide (DMSO) and 1 μl of a tenfold-diluted triethylamine stock solution in DMSO. The reaction was allowed to proceed for 24 h at room temperature. The resulting azide-modified silica particles were then purified by adding 1 ml DI water and centrifuging the particles at 15,000 r.p.m. for 5 min. The supernatant was discarded and the resulting particles were resuspended in 1 ml DI water. This process was repeated seven times, and after the final centrifugation step the particles were resuspended in 100 μl DI water to yield an azide-modified particle stock. The azide-modified particles were stored at 4 °C in the dark and used within 1 month of preparation.

### Synthesis of DNA-functionalized particles

DNA-functionalized particles were created by adding 5 µL of 1mM alkyne-modified DNA solution (“DNA leg”) to a mixture of 5 µL of azide-functionalized silica beads, 5 μl of 2M triethyl ammonium acetate (Glen Research, 60-4110-62), and 25mL of DMSO (MilliporeSigma, MX1457-7). Then, 4 μl of a supersaturated stock solution of ascorbic was added to the reaction to act as a reducing agent. Finally, 2 μl of a 10 mM copper tris[(1-benzyl-1H-1,2,3-triazol-4-yl)methyl]amine (Cu-TBTA, Lumiprobe, 21050) stock solution in 55 vol% DMSO was added to the mixture to initiate copper-catalysed azide–alkyne cycloaddition (CuAAC). The mixture was left to incubate at room temperature for 24 hr. The particles were then purified by washing via centrifugation with the same method described in the synthesis of azide-functionalized silica beads section but with a 10% (w/v) TritonX-100 solution 3 times, and then in 1X PBS 4 times.

### Particle translocation

Before initiating experiments, surfaces presenting DNA substrate were washed with 2 mL of 1X PBS to remove any unbound strands and then with 1X rolling buffer (60 mM NaCl, 20 mM Tris HCl, and 5 mM MgCl₂, pH 7.9). Surface coverage was assessed by fluorescence imaging to ensure homogeneity and adequate signal intensity. DNA-functionalized silica particles were then introduced to the surface by diluting 5 μL of motor particles in 45 μL of 1X PBS. The particles were allowed to hybridize with the complementary surface-bound DNA fuel strands during a 30-minute incubation at room temperature. Following hybridization, unbound particles were removed by gentle washing with 1X PBS. Rolling assays were initiated by replacing the wash buffer with 50 μL of rolling buffer containing 8-16 nM Exonuclease III. Exonuclease III was freshly prepared and added to the rolling buffer immediately before the start of the experiment. Particle movement was monitored using bright-field time lapse imaging at 5 second intervals for 30 minutes using Nikon Elements software. Timelapse and fluorescence image data were then saved for downstream trajectory analysis.

### Particle translocation with RNase H

Experiments were conducted using the same protocol described above in the “Particle Translocation” section, with the following modifications: 5 U of RNase H was added to the surface and imaged for 30 min following initial motor sedimentation. The surface was then washed with 1× rolling buffer, after which the standard protocol was resumed with the addition of Exonuclease III.

### Motion-based viral biosensing

Viral samples were diluted to the desired concentrations, aliquoted, and stored at −80 °C. Prior to biosensing experiments, samples were passively thawed on the benchtop for 5 min and then incubated with aptamer-modified DNA motor silica particles (90% DNA leg + 10% aptamer) in viral matrix for 30 min. The resulting mixture was added onto a surface functionalized with 50% DNA fuel and 50% aptamer strands and allowed to sediment for 30 min. Finally, the viral matrix solution was replaced with rolling buffer containing 16 nM ExoIII, and imaging was carried out following the standard “particle translocation” protocol described above.

### Image processing and particle tracking

Image processing and particle tracking were performed using both Fiji (ImageJ) and custom Python scripts. Nikon Elements image files (*.nd2) were imported into Fiji via the Bio-Formats plugin, where basic pre-processing such as cropping, brightness adjustment, and background subtraction was performed. Motor trajectory analysis, including calculations of net displacement and step size, was carried out in Python (v3.7.4). Drift correction was implemented based on the StackReg^40^ & Fast4DRej^41^ plugins in Fiji. The complete analysis pipeline for extracting quantitative trajectory data from bright-field image sequences was developed collaboratively by Luona Zhang and Yusha Imtiaz. Joshua Hardin developed the computational script for calculating localization errors. Statistical analyses and graphing were performed using GraphPad Prism (v8.4.3).

### Native PAGE

To prepare a 10% native polyacrylamide gel, a solution was made by combining 5 mL of 40% 29:1 acrylamide:bis-acrylamide, 2 mL of 10X TBE buffer, and 14 mL of deionized water in a 50 mL conical tube. The mixture was shaken vigorously until contents were well mixed. Polymerization was initiated by adding 12 μL of TEMED and 60 μL of fresh 30% (w/v) ammonium persulfate (APS), followed by quick vortexing. The solution was poured between 1 mm gel plates (Bio-Rad) and allowed to polymerize for approximately 30 minutes at room temperature. After polymerization, gels were used immediately or wrapped in a damp paper towel and stored at –4°C for up to three days. For gel electrophoresis, DNA oligonucleotide substrates were annealed at 100 nM in a Bio-Rad T100 thermal cycler using stepwise cooling to ensure stable duplex formation. Digestion was initiated by incubating samples at 37°C for 5 minutes with 8 nM Exonuclease III, and then subsequently halted via denaturation by heating the samples to 95°C for 5 mins. A volume of 15 μL from each reaction was mixed with 3 μL of 1X SDS-free Purple DNA loading dye (NEB), which contains 0.042% bromophenol blue and 10% glycerol. Electrophoresis was performed using a Bio-Rad Mini-PROTEAN Tetra Vertical Electrophoresis Cell and PowerPac Basic Power Supply. Gels were pre-run at 80 V for 15 minutes prior to sample loading, and then run at 150 V for 45 minutes. Gels were imaged using an Invitrogen iBright CL750 Imaging System, using Nucleic acid, Cy3, Cy5, or FAM filters to detect labeled strands. Following this, gels were stained with a 1:10,000 dilution of Promega Diamond Nucleic Acid Dye for 10 minutes and re-imaged in nucleic acid mode to visualize total DNA content and assess strand degradation in response to ExoIII treatment.

### Changepoint algorithm development

To identify and quantify bursts in motor velocity, a changepoint detection algorithm was implemented in MATLAB (**Supplementary Fig. 11**). First, the instantaneous velocity of each bead in the x and y-dimensions was calculated and smoothed using a moving average (5 frames). These velocities were then combined to yield a smoothed, total displacement per frame (ΔXY). A changepoint detection algorithm^42^ was then applied to the ΔXY trace to segment each trajectory into periods of statistically distinct mobility, and the probability distribution function of these mobilities was fit to two Gaussians. Lastly, these Gaussian fits were used to compute a threshold to identify high-mobility bursts of motion (where the threshold was set to 3 standard deviations above the mean for the low-mobility Gaussian fit, see **Supplementary Figure 11D**).

## Acknowledgements

We acknowledge support from NIH U01AA029345-01 (K.S.), Merck Future Insight Prize (K.S.), NSF Graduate Research Fellowship Program (K.J.), NIGMS T32GM152344 (J.H.), and Emory University and the Biological Discovery through Chemical Innovation Training Program (J.H.). We thank Sergey Urazhdin for access to the thermal evaporator. We also acknowledge Hiroaki Ogasawara, Tharindu Rajasooriya, and Vageesha Herath for their assistance with preparing and purifying DNA oligonucleotide samples. We thank Dr Richard Webby and Dr. Anami Patel for contributing UV-inactivated SARS-CoV-2 samples. The authors acknowledge the use of a large language model (ChatGPT, OpenAI) to assist in developing custom Python scripts and to provide grammatical and stylistic suggestions during the preparation of this manuscript.

## Author Information

### Contributions

Y.I., A.B., and K.S. conceived the project. Y.I. designed and conducted all the experiments, analyzed data, created computational scripts, and compiled the figures. J.H. helped conduct experiments, analyzed data, and created computational scripts. L.Z. and A.F. helped create computational scripts. B.S., M.R., and K.J. helped conduct experiments. Y.I. and K.S. wrote the manuscript. J.H. and S.P. assisted in writing the manuscript.

## Ethics declarations

Programming scripts used in this publication can be found on the SalaitaLab GitHub page.

